# Mediator is essential for small nuclear and nucleolar RNA transcription in yeast

**DOI:** 10.1101/352906

**Authors:** Jason P. Tourigny, Moustafa M. Saleh, Gabriel E. Zentner

**Affiliations:** Department of Biology, Indiana University, Bloomington, IN, USA

## Abstract

Eukaryotic RNA polymerase II (RNAPII) transcribes mRNA genes as well as non-protein coding RNAs (ncRNAs) including small nuclear and nucleolar RNAs (sn/snoRNAs). In metazoans, RNAPII transcription of sn/snoRNAs is facilitated by a number of specialized complexes, but no such complexes have been discovered in yeast. It has thus been proposed that yeast sn/snoRNA promoters use the same complement of factors as mRNA promoters, but the extent to which key regulators of mRNA genes act at sn/snoRNA genes in yeast is unclear. Here, we investigated a potential role for the Mediator complex, essential for mRNA gene transcription, in the transcription of sn/snoRNA genes. We found that the complete Mediator complex maps to most sn/snoRNA gene regulatory regions and that loss of Mediator function results in a robust reduction in RNAPII and TFIIB occupancy at sn/snoRNA genes. Furthermore, deletion of subunits of the activator-interacting Mediator tail module does not affect Mediator recruitment to, or transcription of, sn/snoRNAs. Taken together, our analyses indicate that Mediator promotes PIC formation and transcription at sn/snoRNA genes, expanding the role of this critical regulator beyond its known functions in mRNA gene transcription and demonstrating further mechanistic similarity between the transcription of mRNA and sn/snoRNA genes.

## Introduction

Eukaryotic RNA polymerase II (RNAPII), responsible for the transcription of mRNA genes, also transcribes several classes of non-protein coding RNA (ncRNA) genes. Two prominent classes of RNAPII-transcribed eukaryotic ncRNA genes are small nuclear and small nucleolar RNAs (sn/snoRNAs). snRNAs are involved in pre-mRNA splicing (1), while snoRNAs primarily function in ribosomal RNA (rRNA) maturation via methylation and pseudouridylation of specific rRNA bases (2).

Most sn/snoRNA transcription in eukaryotes is carried out by RNAPII, with a few notable RNAPIII-transcribed exceptions including the *U6* snRNA gene (3). While transcription of both mRNA and the majority of sn/snoRNA genes is carried out by RNAPII, notable differences between their regulation have been described, particularly in metazoans. In addition to the documented dependence of a handful of sn/snoRNAs on several general transcription factors (GTFs) that are components of the pre-initiation complex and generally essential for RNAPII transcription (TFIIA, TFIIB, TFIID, TFIIE, TFIIF, and TFIIH) (4–6), metazoan sn/snoRNA gene transcription relies on a number of specialized factors. One such factor is the small nuclear RNA activating complex (SNAPc), which contains the TATA-binding protein (TBP) and five specific subunits and is essential for transcription from sn/snoRNA promoters *in vitro* (7–9). Metazoan sn/snoRNA gene transcription also involves the little elongation complex (LEC), which promotes RNAPII occupancy and elongation at these genes (10). Lastly, the Integrator complex, which contributes to both transcriptional elongation and 3’ end processing, is involved in metazoan sn/snoRNA expression (11, 12). Metazoan RNAPII-regulated sn/snoRNA promoters also contain specialized DNA motifs, the distal and proximal sequence elements (DSE and PSE), both of which are also found in type 3 RNAPIII promoters (13). The DSE is similar to an enhancer element, containing binding sites for a number of transcription factors (TFs) including Oct1 and ZNF143 (14). Oct1 and ZNF143 enhance the transcription of sn/snoRNAs by promoting the association of SNAPc with the PSE (7, 14–16). Notably, TATA boxes are present in RNAPIII- but not RNAPII-regulated sn/snoRNA promoters (14), and insertion of a TATA box into the promoter of the RNAPII-transcribed *U2* snRNA gene switches it to a target of RNAPIII (17, 18).

In contrast to metazoans, no sn/snoRNA-specialized transcriptional regulatory complexes or sequence elements have been described in yeast, which uses RNAPII to transcribe all sn/snoRNA genes with the exception of the RNAPIII-transcribed *U6* snRNA and the *snR52* snoRNA (19). Instead, it has been proposed that yeast RNAPII uses the same set of factors to facilitate the transcription of mRNA and sn/snoRNA genes. Indeed, early studies of the promoter region of the polycistronic *snR78-72* gene defined a binding site for the telomere-binding TF Rap1, a key regulator of ribosomal protein gene expression (20), an AT-rich region, and a TATA box (21). A genomic analysis of yeast snoRNAs revealed the presence of these elements in various proportions across 57 promoters as well as additional motifs including binding sites for the TF Reb1, rRNA processing elements, and binding sites for the telomere-associated TF Tbf1, which was reported to activate transcription of the *snR64* snoRNA gene (22). Beyond these observations, however, the extent to which mRNA and sn/snoRNA genes depend on the same RNAPII-associated factors for expression is unclear.

One promising candidate for a regulatory factor used by RNAPII at both mRNA and sn/snoRNA genes is Mediator, a modular, evolutionarily conserved complex required for the majority of mRNA transcription in yeast (23–25). The 25 subunits of yeast Mediator are divided into four modules: head, middle, tail, and kinase. Mediator associates with transcriptional activators at distal regulatory elements via its tail module and RNAPII at promoters via its head module, thus integrating distinct regulatory inputs to promote assembly of the PIC and subsequent transcriptional initiation (26). While the role of Mediator in mRNA transcription has been extensively studied, little is known about its relationship to RNAPII transcription of ncRNAs. In mouse embryonic stem cells, Mediator forms a meta-coactivator complex (MECO) with the Ada-Two-A-containing (ATAC) histone acetyltransferase complex that associates with a small number of snRNA genes to promote their expression (27). Furthermore, the metazoan-specific Mediator subunit Med26 has been implicated in the transcription of a small number of sn/snoRNA genes in mouse and human cells via recruitment of LEC (28). These observations suggest that Mediator is an important regulator of sn/snoRNA gene transcription; however, important questions remain. First, as the above studies analyzed a selected few loci, it is unclear if any role of Mediator in promoting sn/snoRNA gene transcription is global. Moreover, as the above studies analyzed Mediator function in the context of interactions with metazoan-specific complexes (ATAC, LEC) and a metazoan-specific Mediator subunit (Med26), it is unknown if Mediator functions in organisms lacking these components of the transcription machinery. Lastly, it has not been tested if the function of Mediator in PIC assembly is relevant at sn/snoRNA gene promoters.

Here, we sought to determine if Mediator plays a global role in the transcription of sn/snoRNA genes in yeast. Using genome-wide analyses, we found that upstream regions potentially equivalent to mRNA gene upstream activating sequences (UASs) and promoters of sn/snoRNA genes are occupied by Mediator. Inducible depletion of the structurally essential Mediator subunit Med14 results in equivalent reductions in RNAPII association with sn/snoRNA and mRNA genes, indicating an essential role of the complete Mediator complex in sn/snoRNA gene transcription. Similar to its function at mRNA genes, Mediator promotes PIC assembly at sn/snoRNA gene promoters. Interestingly, removal of subunits of the Mediator tail module, responsible for interactions with transcriptional activators, does not affect Mediator recruitment to sn/snoRNA genes. Our results indicate that tail-independent recruitment of Mediator to sn/snoRNA genes is required for their transcription and suggest mechanistic similarities in Mediator activity at yeast mRNA and sn/snoRNA genes.

## Results

### Mediator associates with sn/snoRNA genes

We first assessed the genomic localization of Mediator with respect to sn/snoRNA genes using chromatin endogenous cleavage and high-throughput sequencing (ChEC-seq) (29), which we previously showed effectively maps Mediator binding to UASs (30). However, we only mapped subunits of the Mediator head module. In order to interrogate the localization of the entire Mediator complex, we mapped the Med8 subunit from the head module as well as the scaffold subunit Med14 and the tail module subunit Med3. Visual inspection of ChEC-seq and ChIP-seq data revealed enrichment of Med8, Med14, and Med3 at the intergenic region of the divergent *snR31* and *snR5* snoRNA genes (Fig. 1A). To analyze these data more systematically, we compiled a list of 70 RNAPII-transcribed sn/snoRNA genes not overlapping protein-coding genes, condensed to 58 loci due to the existence of five polycistronic snoRNA clusters in the yeast genome (31) (see Materials and Methods), and aggregated data at these positions. This analysis revealed strong peaks of Med8, Med14, and Med3 enrichment over free MNase signal upstream of sn/snoRNA TSSs (Fig. 1B). Notably, the position of maximum Mediator subunit occupancy was relatively far upstream of sn/snoRNA TSSs (421 bp for Med8 and 403 bp for Med14 and Med3), potentially consistent with Mediator association with upstream activating sequences (UASs), where it binds most prominently to mRNA genes under normal conditions (30, 32). Notably, the dynamic ranges and signal-to-noise ratios of the Med14 and Med3 ChEC-seq data were much higher than that of the Med8 data at sn/snoRNA genes. This may reflect the potentially shorter distances of the C-termini of Med14 and Med3 than Med8 to DNA (33). To more quantitatively assess enrichment of Mediator at sn/snoRNA genes, we called peaks on the Med14 ChEC-seq dataset with the free MNase dataset as a control. We then assessed the 58 analyzed sn/snoRNA TSSs for proximal peaks. Of these, 41/58 (70.7%) displayed a significant Med14 peak at the threshold used, indicating that the majority of sn/snoRNA promoters.

**Figure 1.**
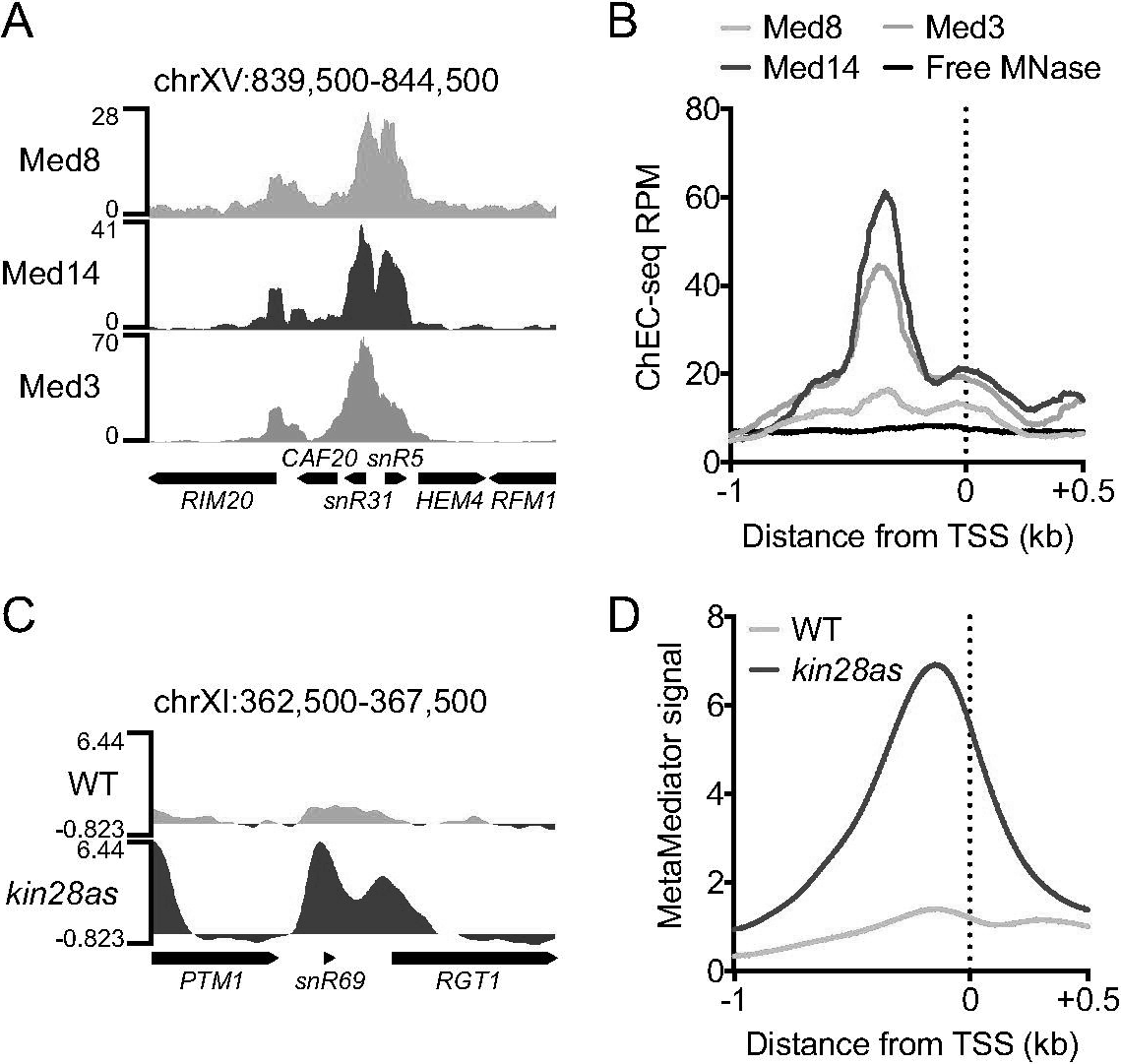
Mediator associates with sn/snoRNA genes. (A) Genome browser view of Med8, Med14, and Med3 ChEC-seq signal at a representative sn/snoRNA-containing region of the yeast genome. (B) Average plot of Med8, Med14, Med3, and free MNase ChEC-seq signal at sn/snoRNA genes (n = 58). (C) Genome browser view of MetaMediator ChIP-chip signal in 1-NA-PP1-treated WT and *kin28as* cells at a representative sn/snoRNA-containing region of the yeast genome. (D) Average plot of MetaMediator ChIP-chip signal in 1-NA-PP1-treated WT and *kin28as* cells at sn/snoRNA genes.

Having observed association of complete Mediator with putative sn/snoRNA gene UASs, we sought to determine if we could also detect Mediator association with sn/snoRNA gene promoters, which occurs via interactions of the head module with the carboxy-terminal domain (CTD) of RNAPII (34). When the CTD is phosphorylated by the TFIIH kinase Kin28, Mediator is rapidly released from the PIC (35), and it has been shown that efficient detection of promoter-associated Mediator by ChIP requires impairment of Kin28 function in order to trap Mediator in complex with the PIC (32, 36). We previously found that ChEC-seq does not map Mediator binding to promoters upon Kin28 inhibition, presumably due to occlusion of promoter DNA by the PIC (30). Thus, to interrogate Mediator association with sn/snoRNA promoters, we obtained ChIP-chip data from a previous study in which the genome-wide association of twelve Mediator subunits (“MetaMediator”) was profiled following inhibition of *kin28as*, a form of Kin28 sensitive to the ATP analog 1-Naphthyl PP1 (1-NA-PP1) (33). Visual inspection of MetaMediator signal at the *snR69* snoRNA gene revealed little upstream enrichment of Mediator in wild-type (WT) cells treated with 1-NA-PP1 (Fig. 1C), consistent with the reported low ChIP efficiency of Mediator at the upstream regions of many highly-transcribed genes (32, 37). However, *kin28as* cells treated with 1-NA-PP1 displayed a marked increase in MetaMediator enrichment just upstream of the *snR69* TSS (Fig. 1C). Systematic analysis of MetaMediator occupancy in WT and *kin28as* cells confirmed this single-locus observation across the genome (Fig. 1D). Based on these observations, we conclude that the complete Mediator complex associates with both putative UASs and promoters at sn/snoRNA genes, suggesting mechanistic similarities in the activity of Mediator at these distinct classes of genes.

### RNAPII occupancy of sn/snoRNA genes is Mediator-dependent

Having established that most sn/snoRNA genes are bound by Mediator, we set out to determine what effect its loss would have on sn/snoRNA gene transcription. We used native RNAPII ChIP-seq data from a previous study in which Med14, essential for the structural integrity of Mediator (23, 38), was depleted from yeast cells using the auxin degron system (24). In this approach, a target protein is tagged with an auxin-inducible degron (AID) consisting of a 3xV5 tag and the auxin repressor protein IAA7 in a yeast strain constitutively expressing the *Oryza sativa* ubiquitin ligase OsTIR1. In the presence of auxin, a complex is formed between *Os*TIR1 and the AID-tagged protein, resulting in ubiquitylation and degradation of the target protein (39). This method was reported to yield nearly complete destruction of Med14 within 30 min (24). Importantly, the reported ChIP-seq experiments used a defined amount of *S. pombe* cells as a spike-in, allowing for quantification of global changes in RNAPII binding to the genome. Indeed, using this approach, the authors found that Mediator destabilization via Med14 degradation reduced the transcription of nearly all ~4,800 analyzed mRNA genes (24).

We obtained data for two RNAPII ChIP-seq replicates treated with either vehicle (DMSO) or 500 μM of the auxin indole-3-acetic acid (3-IAA) to deplete Med14 as well as replicates of RNAPII ChIP-seq from WT cells treated with DMSO or 3-IAA. We first visualized RNAPII signal at three regions of the genome harboring one or more sn/snoRNAs. WT cells (that is, those bearing *Os*TIR1 but not Med14-AID) displayed robust RNAPII occupancy of the snoRNA genes *snR17b*, *snR190-snR128*, *NME1*, and *snR66* when treated with either DMSO or 3-IAA, as did Med14-AID cells treated with DMSO. However, RNAPII signal at these genes was almost completely eliminated in Med14-AID cells treated with 3-IAA (Fig. 2). We next quantified RNAPII ChIP-seq signal within the 83 bp downstream of each sn/snoRNA TSS (corresponding to the length of the shortest analyzed gene). As observed at the individual genomic regions analyzed above, 3-IAA treatment of WT yeast had essentially no effect on RNAPII occupancy of either sn/snoRNA or mRNA genes, for which we quantified RNAPII ChIP-seq signal in the 100 bp downstream of each TSS (Fig. 3). However, depletion of Med14 strongly reduced RNAPII ChIP-seq signal in all 58 tested sn/snoRNA genes (median log_2_(fold change) = −3.74), and the extent of the reduction in sn/snoRNA gene RNAPII occupancy was comparable to that observed for the 1,000 most highly-RNAPII occupied mRNA in DMSO-treated Med14-AID cells (median log_2_(fold change) = −3.62) (Fig. 3).

**Figure 2.**
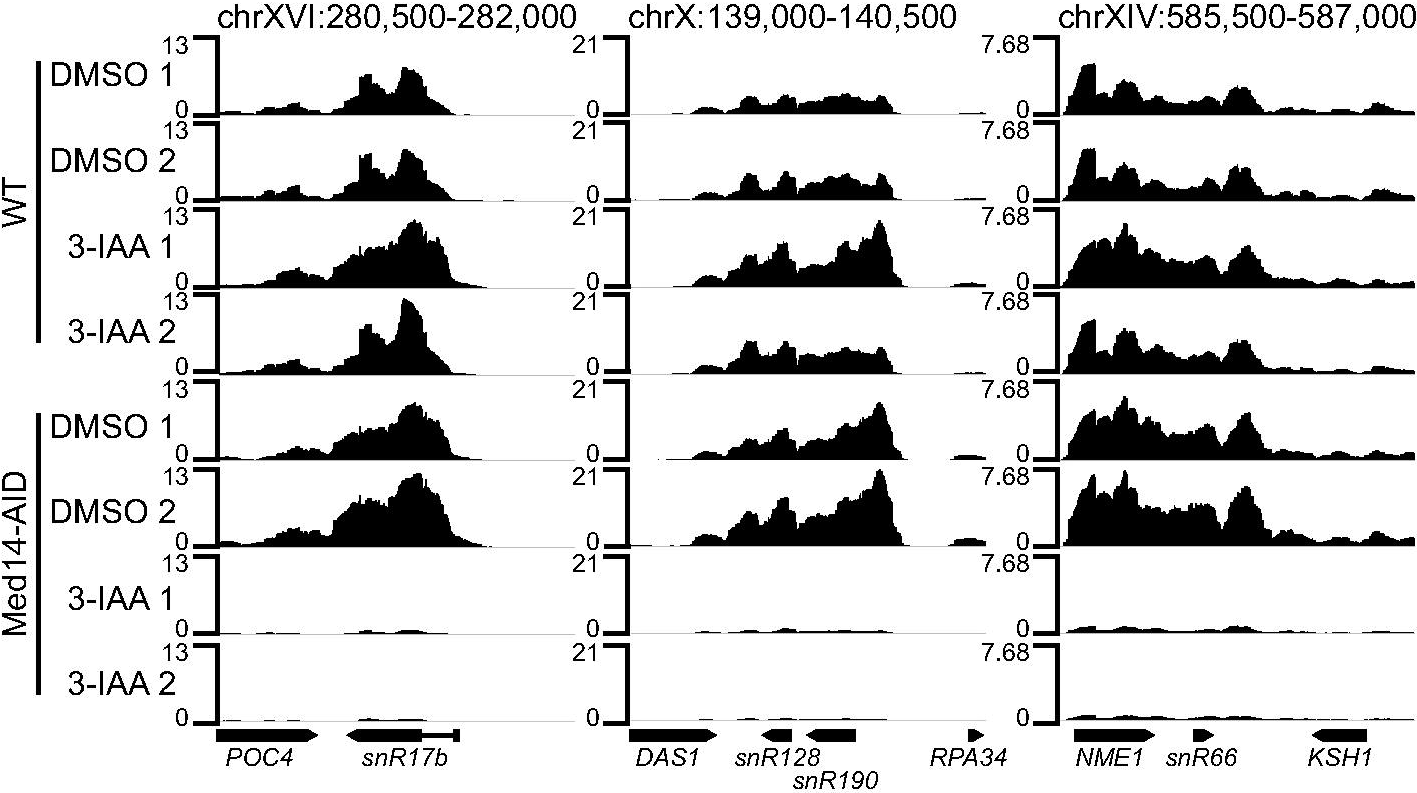
Mediator depletion reduces RNAPII binding to sn/snoRNA genes. Genome browser views of spike in-normalized RNAPII ChIP-seq signal from WT and Med14-AID strains at three regions of the yeast genome containing one or more sn/snoRNA genes.

**Figure 3.**
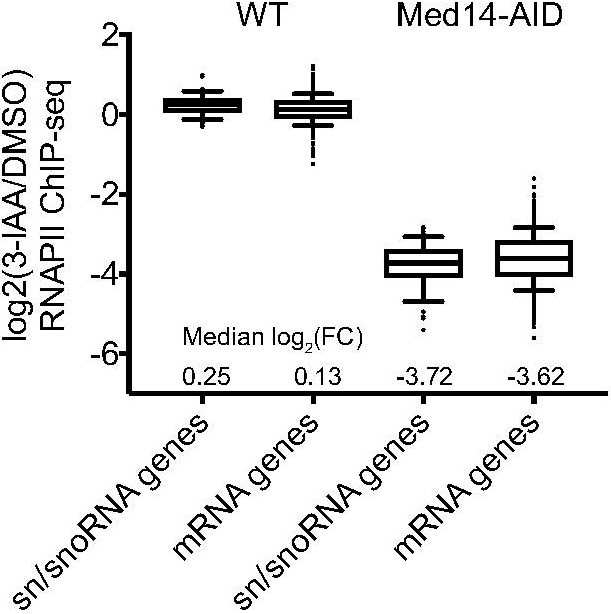
Mediator loss globally reduces RNAPII occupancy in sn/snoRNA genes. Boxplots of spike in-normalized log_2_(3-IAA/DMSO) RNAPII ChIP-seq signal at sn/snoRNA and mRNA genes in WT and Med14-AID strains. The median log_2_(fold change) in RNAPII ChIP-seq signal for each condition is provided.

### Mediator promotes PIC formation at sn/snoRNA gene promoters

A major function of Mediator *in vitro* and *in vivo* is proposed to be stimulation of PIC formation (32, 40–43). Given that loss of Mediator results in a marked reduction in RNAPII occupancy within sn/snoRNA gene coding regions (Fig. 2-3), we asked if it would also reduce PIC formation at sn/snoRNA gene promoters. To this end, we constructed a Med14-AID strain with a FLAG-tagged form of *SUA7*, encoding TFIIB, to allow ChIP-seq analysis of PIC formation. Consistent with previous data (24), we observed near-total degradation of Med14 within 30 min of 3-IAA addition (Fig. 4A). In line with the essential role of Med14, Med14-AID cells did not grow on solid medium containing 3-IAA (Fig. 4B). To measure PIC aseembly at sn/snoRNA gene promoters, we performed two replicates of TFIIB ChIP-seq following a 30 m treatment with DMSO or 3-IAA. Consistent with the reduction in RNAPII occupancy we observed above (Fig. 2-3), most analyzed sn/snoRNA promoters displayed moderately reduced TFIIB binding (median log_2_(fold change) = −0.95), similar to what we observed at the 1,000 most highly TFIIB-occupied mRNA gene promoters (median log_2_(fold change) = −0.88) (Fig. 4C). This observation indicates that the role of Mediator in promoting PIC formation is conserved between mRNA and sn/snoRNA genes.

**Figure 4.**
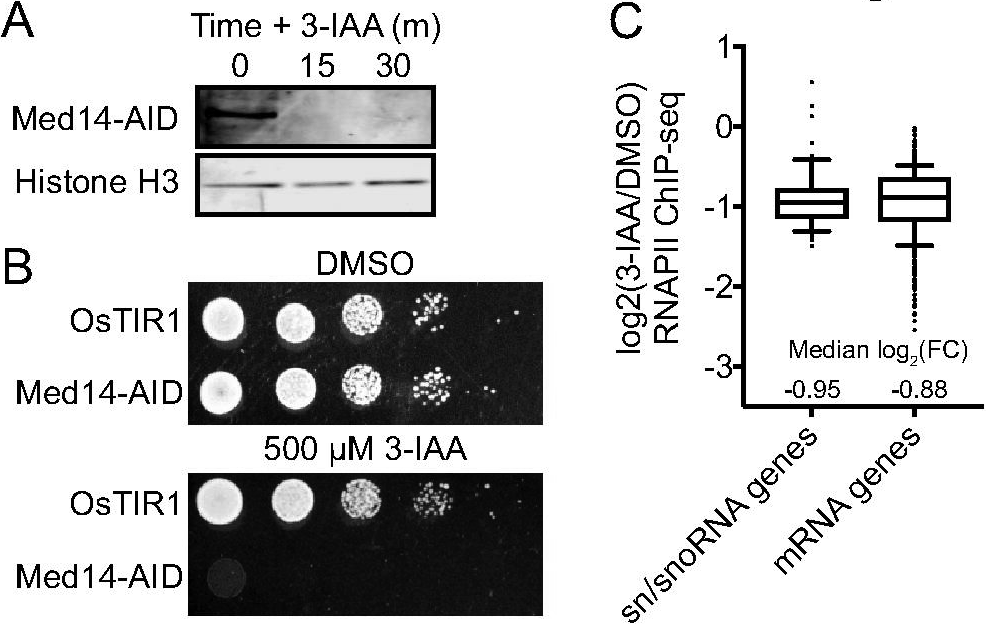
Mediator destabilization reduces PIC formation at sn/snoRNA promoters. (A) Western blot showing the kinetics of Med14-AID depletion. A Histone H3 blot is shown as a loading control. (B) Spot assays analyzing the growth of parental and Med14-AID strains on YPD containing DMSO or 500 μM 3-IAA. (C) Boxplots of RPM-normalized log_2_(3-IAA/DMSO) TFIIB ChIP-seq signal at sn/snoRNA and mRNA genes in WT and Med14-AID strains.

### The Mediator tail module is not required for sn/snoRNA transcription

At mRNA genes, Mediator binds most prominently to UASs via interactions with transcriptional activaors and only transiently associates with core promoters (32, 36). We thus asked if the Mediator tail, responsible for activator interactions, is required for recruitment of Mediator to sn/snoRNA genes. We tested if deletion of Med15, a tail subunit and major target of activators (44, 45), would impact the recruitment of Mediator to sn/snoRNA genes. We queried our previous ChEC-seq data in which binding of Med17 was mapped in either a WT or *med15*Δ background (30), assessing the effects of *MED15* deletion on Mediator binding to sn/snoRNA genes and mRNA genes classified by SAGA/TFIID coactivator dependence (46), as the Mediator tail module has been reported to act most prominently at SAGA-dominated genes (47). Consistent with the reported dependence of SAGA-dominated genes on the Mediator tail module, Med17 binding at the UASs of the 50 most Mediator-enriched SAGA-dominated gene UASs was markedly decreased in the *med15*Δ strain (median log_2_(fold change) = −2.59), while Med17 enrichment at the 500 most Mediator-enriched TFIID-dependent gene UASs was on average only slightly decreased (median log_2_(fold change) = −0.39) (Fig. 5A). In contrast, Med17 association with sn/snoRNA gene upstream regions was slightly increased in the *med15*Δ background (median log_2_(fold change) = 0.52) (Fig. 5A), indicating that activator-tail module interactions may not be a major determinant of Mediator recruitment to sn/snoRNAs in yeast.

**Figure 5.**
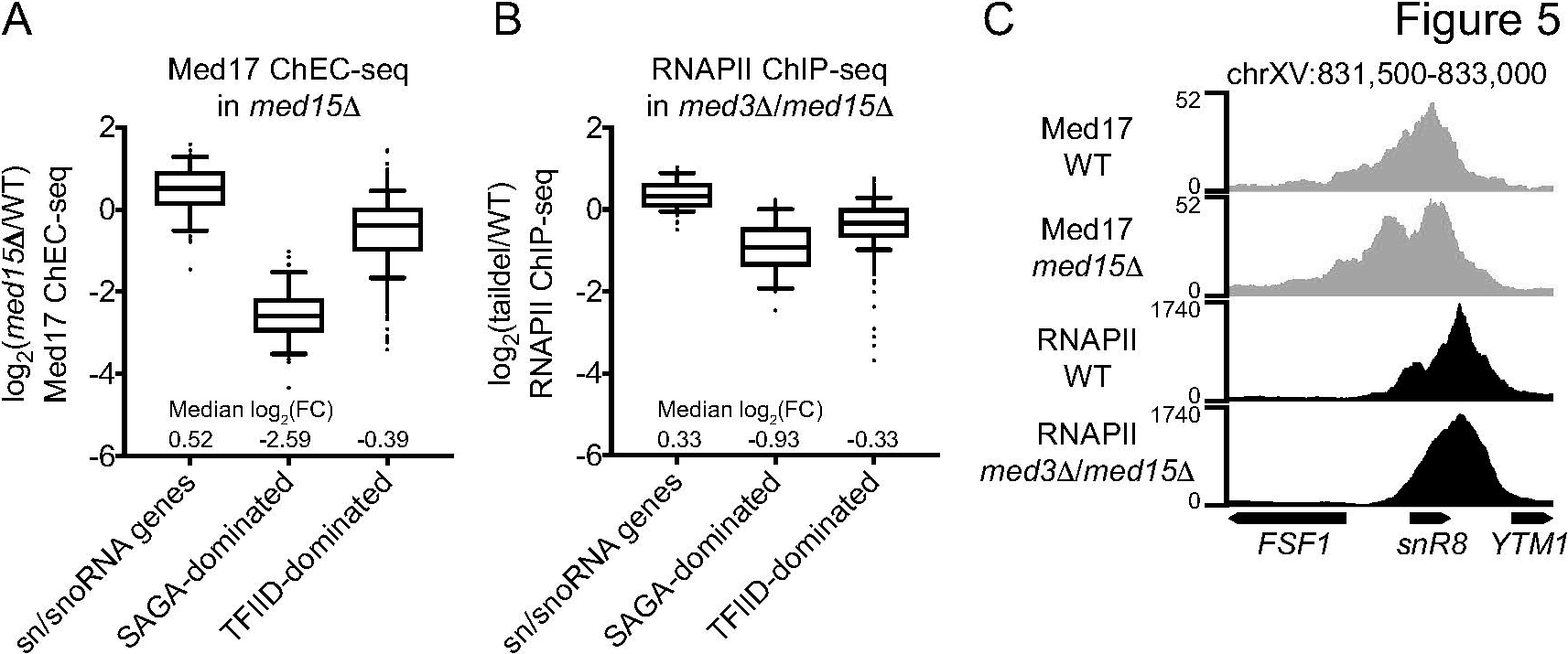
Mediator tail depletion does not affect Mediator association with sn/snoRNA genes or sn/snoRNA transcription. (A) Boxplots of RPM-normalized log_2_(*med15*D/WT) Med17 ChEC-seq signal at sn/snoRNA genes, the top 50 most highly Med17-occupied SAGA-dominated genes, and the top 500 most highly Med17-occupied TFIID-dominated genes. The median log_2_(fold change) in Med17 ChEC-seq signal for each condition is provided. (B) Boxplots of RPM-normalized log_2_(taildel/WT) RNAPII ChIP-seq signal at sn/snoRNA genes, the top 50 most highly Med17-occupied SAGA-dominated genes, and the top 500 most highly Med17-occupied TFIID-dominated genes. The median log_2_(fold change) in Med17 ChEC-seq signal for each condition is provided. (C) Genome browser views of Med8 ChEC-seq signal in WT and *med15*Δ yeast and RNAPII ChIP-seq signal in WT and tail deletion yeast at the *snR8* locus.

To investigate the potential dispensability of the Mediator tail for sn/snoRNA transcription, we analyzed RNAPII ChIP-seq data from WT yeast and a strain lacking *MED3* and *MED15*, hereafter referred to as “tail deletion” (48). In line with the aforementioned dependence of SAGA-dominated genes on the tail, the 50 most highly RNAPII-bound SAGA-dominated genes displayed a strong decrease in RNAPII ChIP-seq signal in the tail deletion strain (median log_2_(fold change) = −0.93), while the top 500 most robustly RNAPII-enriched, TFIID-dominated genes were substantially less affected (median log_2_(fold change) = −0.33) (Fig. 5B). In contrast, RNAPII ChIP-seq signal at sn/snoRNA genes was essentially unaffected in the tail deletion strain (median log_2_(fold change) = 0.33) (Fig. 5B). Visual inspection of the *snR8* locus confirmed the lack of effect of tail subunit deletion on Mediator and RNAPII binding (Fig. 5C). Taken together, these observations strongly suggest that the tail module is dispensable for Mediator regulation of sn/snoRNA transcription.

## Discussion

Mediator is a conserved, essential transcriptional regulatory complex recently shown to be essential for the majority of RNAPII mRNA transcription in yeast. Here, we investigated its contribution to the transcription of sn/snoRNA genes in a tail-independent manner and promotes their transcription at least in part by facilitating PIC assembly.

Our analysis of Mediator binding to the genome suggests that the regulatory organization of sn/snoRNA gene upstream regions is similar to that of mRNA genes. ChEC-seq, which we previously showed efficiently maps Mediator to UASs (30), delineated a peak of Mediator occupancy ~400 bp upstream of sn/snoRNA gene TSSs on average. This position is consistent with the location of UASs at mRNA genes (49), though this distance measurement may be an over- or underestimate, as MNase cleavage in ChEC occurs upstream or downstream of the tagged protein depending on its orientation. We also found that Mediator could be robustly detected at sn/snoRNA promoters upon inhibition of the TFIIH kinase Kin28, consistent with previous observations at mRNA promoters (32, 36). This observation is also in line with the previously described role of Cdk7, the human ortholog of Kin28, in promoting RNAPII CTD phosphorylation at the *U1* and *U2* snRNA genes in human cells (5). This apparent organizational conservation of Mediator binding to sn/snoRNA and mRNA gene upstream regions is in line with the similar effects of Mediator depletion on RNAPII and TFIIB occupancy observed among both gene classes.

Our results also bear on how Mediator is recruited to sn/snoRNA genes. We found that deletion of the major activator-binding tail subunit Med15 has little effect on Mediator association with sn/snoRNA genes and that combined deletion of the tail subunits Med3 and Med15 does not reduce RNAPII occupancy of sn/snoRNA genes. These observations may indicate that, in terms of Mediator recruitment and function, sn/snoRNA genes are similar to genes traditionally classified as TFIID-dominated, whose expression is generally insensitive to loss of Mediator tail function (47). The tail has also been reported to be dispensible for Mediator occupancy at TFIID-dominated genes (48), though this has recently been questioned (33). One possible mechanism for recruitment of Mediator to sn/snoRNAs is via the interaction of activators, including Tbf1, with other subunits of Mediator such as the tail/middle module connector Med16, which interacts with activators such as Gcn4 in yeast (45), DIF in *Drosophila* (50), and NRF2 in mouse and human (51). Indeed, a recent proteomic analysis of Mediator interactions reported a modest interaction between Mediator and Tbf1 (52). Mediator could also be recruited to sn/snoRNA genes independent of activators via interactions with the PIC (33, 53) (bioRxiv: https://doi.org/10.1101/207282).

Taken together, our results indicate that Mediator is globally required for sn/snoRNA transcription in yeast and that its core function, the facilitation of PIC formation, is conserved between mRNA and sn/snoRNA genes. Moving forward, it will be of interest to determine via genetic studies if the Mediator-bound upstream regions of sn/snoRNA genes enhance their transcription and are thus analogous to mRNA gene UASs. Studies to delineate the mechanisms of recruitment of Mediator to sn/snoRNA genes will also provide further insight into the degree of commonality in Mediator function and regulation between mRNA and sn/snoRNA genes.

## Materials and Methods

### Yeast Methods

Med8, Med14, and Med3 were tagged with 3xFLAG-MNase-HIS3MX6 using pGZ109 (30). The auxin degron parental strain GZY191 was generated by transformation of pSB2273 (kindly provided by Matthew Miller), encoding *GPD1* promoter-driven *Os*TIR1, into the *HIS3* locus in a wild-type BY4705 strain. *MED14* was tagged with AID using pL260 (kindly provided by Matthew Miller), encoding 3xV5-IAA7-kanMX6, resulting in strain GZY219. Lastly, *SUA7* was tagged with 3xFLAG using pFA6a-6xGLY-3xFLAG-hphMX4 (a gift from Mark Hochstrasser, Addgene plasmid #20755), resulting in strain GZY231. Strain details are given in Table 1. Cells were grown in YPD at 30°C under constant agitation. For auxin treatment, a 100 mL culture at an OD_600_ of >0.8 was split into two flasks. One culture was treated with DMSO, the other with 500 μM 3-IAA dissolved in DMSO, for 30 min.

### ChEC-seq

ChEC-seq was performed as described (29) except for the size selection step. RNase-treated ChEC DNA was brought up to 200 μL with 10 mM Tris pH 8.0 and 160 μL SPRI beads (54) were added (0.8:1 beads:sample ratio versus the 2.5:1 ratio used in the original protocol). The sample was pipetted up and down ten times to mix and incubated at RT for 5 min. Beads were collected on a magnetic rack for 2 min and the supernatant was collected for DNA isolation. Sequencing libraries were prepared at the Indiana University Center for Genomics and Bioinformatics (CGB) using the NEBNext Ultra II Library Prep Kit for Illumina. Libraries were sequenced in paired-end mode on the Illumina NextSeq 500 platform at the CGB. Read lengths were 79 bp for Med8 and 42 bp for Med14 and Med3.

### ChIP-seq

After DMSO or 3-IAA treatment, cultures were fixed with formaldehyde (1% final) for 10 minutes at room temperature (RT), then quenched with glycine (125 mM final) for 5 minutes. After washing with PBS, fixed cell pellets were preserved at −80°C. Thawed pellets were later spheroplasted by resuspension in PBS along with 90 μl of 5 mg/mL zymolyase for 5 m at 37°C. All ChIP solutions henceforth were supplemented with a protease inhibitor cocktail. After gentle centrifugation and washing with 1ml PBS, ~100ul spheroplasts were lysed with the addition of 50μL lysis buffer (50 mM Tris pH 8.0, 150 mM NaCl, 10 mM EDTA, 3% SDS) at RT for 5 m. Lysates were then diluted to 1.5 mL with 1350 μL of dilution buffer (20 mM Tris pH 8.0, 150 mM NaCl, 2 mM EDTA, 1% Triton X-100) and transferred to a Covaris milliTube on ice. Sonication of diluted ChIP lysate was performed in a Covaris S220 sonicator, with parameters of 150 W Peak Power, 30% Duty Cycle, and 200 Cycles/burst, for a total sonication time of 150 s (on 30s -- cool 15s, times 5). Thereafter, samples were centrifuged for 10 m at 21,000 × *g*and 4°C. The soluble lysates were then transferred to new tubes along with ~30 μL of Sigma FLAG beads (M8823) pre-blocked with BSA, and the IP was performed for 3h at 4°C. Beads were then washed 2 × 1 mL in fresh cold dilution buffer, and 1 × 1ml in LiCl wash buffer (20 mM Tris pH 8.0, 250 mM LiCl, 1 mM EDTA, 1% Triton, 0.1% Nonidet-P40) at RT, with each wash lasting 3 m. For extraction and de-crosslinking, washed beads were resuspended in 250 μL elution buffer (50 mM Tris pH 8.0, 250 mM NaCl, 10 mM EDTA, 1% SDS) with 60 μg proteinase K and incubated overnight at 1,200 RPM and 65°C in an Eppendorf ThermoMixer. The next morning, ChIP DNA was extracted with 200 μL phenol/chloroform/isoamyl alcohol, treated with 10 μg RNase A for 20 m at 37°C, re-extracted with phenol/chloroform, and precipitated with linear acrylamide in >80% EtOH at −80C overnight. Precipitated DNA was pelleted for 20 m at 4°C and 21,000 × *g*, washed with RT 75% ethanol, re-spun for 5 m, dried at RT, and finally resuspended in 20 μL T low-E (50 mM Tris, 0.1 mM EDTA). ChIP-seq libraries were prepared and sequenced as described above for ChEC-seq libraries. Read lengths were 42 bp for the first pair of DMSO/3-IAA replicates and 38 bp for for the second replicate pair.

### Data analysis

#### sn/snoRNA gene list

We used the YeastMine QueryBuilder (https://yeastmine.yeastgenome.org/) to generate a list of sn/snoRNA genes. We first retrieved lists of items with data type “snRNA” (6 items) or “snoRNA” (77 items). For snRNA genes, the RNAPIII-transcribed *snR6* gene was removed and the overlapping *snR7-L* and *snR7-S* genes were condensed into the long isoform. For snoRNA genes, we used the overlapping features function of YeastMine to identify snoRNAs overlapping ORFs (13/77). Two snoRNA genes, *snR43* and *snR73*, were found to overlap a deleted and dubious ORF, respectively, and so were retained in the final list. The RNAPIII-transcribed *snR52* gene was removed from the final list. We also included the RNAPII-transcribed *TLC1* gene, encoding telomerase RNA. This resulted in a list (Dataset S1, tab 1) containing 70 genes (4 snRNAs, 65 snoRNAs, and *TLC1*). For the purposes of our TSS-based analyses, polycistronic snoRNA clusters (*snR190-snR128*, *snR41-snR70-snR51*, *snR67-snR53*, *snR57-snR55-snR61*, and *snR78-snR77-snR76-snR75-snR74-snR73-snR72*) were considered as single genes, with the start coordinate of the first snoRNA in the cluster taken to be the cluster’s TSS. This condensation resulted in a list of 58 loci (Dataset S1, tab 2).

#### ChEC-seq

Paired-end Med8, Med14, and Med3 ChEC-seq and 1 min *REB1* promoter-driven free MNase data (SRR1947784) were aligned to the sacCer3 genome build with Bowtie2 (55) using default settings plus `-I 10 -X 700 --no-unal --dovetail --no-discordant --no-mixed`. The above free MNase dataset was used because it was generated in the W1588-4C background, which is congenic to the W303 background in which the Mediator ChEC-seq datasets were generated except that a weak *RAD5* mutation is corrected (56). Alignment SAM files were used to make tag directories with HOMER (http://homer.ucsd.edu) (57). HOMER `annotatePeaks.pl` was used to average signal. Average plots were generated with GraphPad Prism 7. Med14 peaks were called using HOMER `findPeaks` with the flags `-style factor -F 2 -L 0 -fdr 0.05` with the free MNase dataset as control. These parameters require that a peak be enriched at least twofold over free MNase at an FDR of 0.05 with no requirement for enrichment over local background. The distance of the Med14 peak midpoint closest to each sn/snoRNA TSS was determined with BEDTools `closest` (58) and each peak was visually inspected for its orientation relative to the corresponding sn/snoRNA gene. We required that a peak midpoint be within or no more than 500 bp upstream of an sn/snoRNA gene to be considered associated with that gene. For analysis of Med17 binding, Mediator signal was quantified from −500 to −100 bp relative to the TSS using HOMER `annotatePeaks.pl`. Lists of SAGA- and TFIID-dependent genes were derived from our previous work (30), and we quantified signal at the the 50 most highly Mediator-occupied SAGA-dominated gene UASs and the 500 most highly Mediator-occupied TFIID-dominated gene UASs as determined by signal in the WT strain.

#### ChIP

MetaMediator bedGraph files representing average ChIP-chip signal for 12 Mediator subunits were obtained from the supplementary material of Jeronimo *et al* (33) and visualized as described for ChEC-seq data. Paired-end spike-in RNAPII ChIP-seq data (Rpb3-3xFLAG) (24) were obtained from GEO (GSE97081) and aligned to both the sacCer3 and EF2 (*S. pombe)* genome builds with Bowtie2 as for ChEC-seq. Tag directories were then created with HOMER. For genome browser visualization, we downloaded spike in-normalized wig files from GEO; these tracks thus offer independent confirmation of our systematic analyses of these data. For quantification of sn/snoRNA transcription, the total raw RNAPII signal in an 83 bp window downstream of the TSS (corresponding to length of the shortest analyzed monocistronic snoRNA gene, *snR79*) was determined using HOMER `annotatePeaks.pl`. The obtained values were then multiplied by a spike-in normalization factor N, the log_2_(3-IAA/DMSO) ratio for each gene in each replicate was calculated, and ratios were averaged. For comparison, we analyzed the 1,000 most highly transcribed mRNA genes as determined by average spike in-normalized signal in the two DMSO-treated replicates of each auxin depletion experiment using a 100 bp window downstream of the TSS. The spike-in normalization factor N was calculated as 10,000 / number of reads mapped to the *S. pombe* genome and used by HOMER for generation of a tag directory. Paired-end TFIIB ChIP-seq data were processed as described for ChEC-seq data. TFIIB signal was quantified from −150 to +50 relative to the TSSs of sn/snoRNA genes and the 1,000 most TFIIB-occupied mRNA promoters as determined by average RPM-normalized signal in the −150 to +50 window, the log_2_(3-IAA/DMSO) ratio for each gene in each replicate was calculated, and ratios were averaged. Single-end RNAPII ChIP-seq data (Rpb1) from WT and tail deletion strains (48) were obtained from the SRA (SRP047524) and aligned to sacCer3 with Bowtie2 using default parameters plus `--no-unal`. Tag directories were created and RNAPII signal was quantified as for spike-in RNAPII ChIP-seq data, except that RPM normalization was used. We quantified signal at the 50 most highly RNAPII-occupied, SAGA-dominated genes and the 500 most highly RNAPII-occupied TFIID-dominated genes as determined by signal in the WT strain.

## Data availability

Sequencing data have been deposited with the Gene Expression Omnibus (GSE112721).

## Acknowledgements

We thank Logan Hille for strain construction, Robert Policastro for helpful discussions throughout the course of this work, Sebastian Grünberg for critical reading of the manuscript, and the Indiana University Center for Genomics and Bioinformatics for library construction and sequencing. This work was supported by Indiana University startup funds (to GEZ)

